# Fruit presence induces polar auxin transport in citrus and olive stem and represses hormone release from the bud

**DOI:** 10.1101/2020.07.15.203927

**Authors:** Dor Haim, Liron Shalom, Yasmin Simhon, Lyudmila Shlizerman, Itzhak Kamara, Michael Morozov, Alfonso Ant Albacete Moreo, Rosa M Rivero, Avi Sadka

**Author notes:** Dor Haim, liron shalom, Yasmin Simhon, Lyudmila Shlizerman, Itzahk Kamara, Michael Morozov, Rosa M Rivero, Alfonso Ant Albacete Moreno.

## Abstract

In many fruit trees, heavy fruit load in one year reduces flowering in the following year, creating a biennial fluctuation in yield termed alternate bearing (AB). In subtropical trees, where flowering induction is mostly governed by the accumulation of cold hours, fruit load is thought to generate a signal (AB signal) that blocks the perception of the cold induction. Fruit removal during a heavy-fruit-load year (On-Crop) is effective at inducing flowering only if performed one to a few months prior to onset of the flowering-induction period. We previously showed that following fruit removal, content of the auxin indoleacetic acid (IAA) in citrus buds is reduced, suggesting that the hormone plays a role in the AB signal. Here, we demonstrate that fruit presence generates relatively strong polar auxin transport (PAT) in citrus and olive stems. Upon fruit removal, PAT is reduced and allows auxin release from the bud. Furthermore, using immunolocalization, hormone and gene expression analyses, we show that in citrus, IAA level in the bud and, specifically, in the apical meristem is reduced upon fruit removal. Overall, our data provide support for the notion that fruit presence generates an auxin signal in the bud which may affect flowering induction.

**HIGHLIGHTS:** Heavy fruit load can reduce flowering intensity the following year. Fruit presence, inducing polar auxin transport in stems and inhibiting auxin release from buds, may be a fruit-load signal.

## Introduction

Auxin plays a major role in many aspects of plant development, including seed sprouting, organ morphogenesis, and plant architecture (Reinhardt *et al.*, 2000; Carraro *et al.*, 2006; Shani *et al.*, 2006; Dubrovsky *et al.*, 2008; Liu *et al.*, 2013). It also contributes to the regulation of reproductive processes, ovary-to-seed transition, and fruit set (Aloni *et al.*, 2006; Obroucheva, 2014). Apical dominance is another aspect thought to be partially controlled by auxin; auxin production occurs in the shoot apical meristem (SAM) and in the young leaves, before it is transported toward the roots by polar movement through the stem (Balla *et al.*, 2011, 2016). Axillary buds located along the stem tend to stay dormant until they become far enough from the SAM, or until the SAM is removed. It is thought that the strength of the auxin flow from the bud plays a role as a signal in sink–source interactions, and determines the new dominant bud(s). According to the auxin transport auto-inhibition (ATA) theory, dominance is not always apical; it can also be exerted between sinks of various strengths regardless of their relative positions (primegenic dominance) (Bangerth, 1989). Auxin is produced by various biosynthetic pathways, with tryptophan as the main precursor, although there may be a tryptophan-independent pathway as well (Wang *et al.*, 2015; Di *et al.*, 2016). Polar auxin transport (PAT) distributes the hormone differentially in various plant organs, forming local auxin maxima or minima at the tissue level, or even in a few cells, thus affecting their development and growth (Petrášek and Friml, 2009). The direction of PAT is mostly controlled by AUX/LAX or PIN-FORMED (PIN) family proteins, which are located throughout the cell’s inner and outer membrane in a polar direction. Whereas AUX/LAX proteins are responsible for the influx of auxin into the cell, PIN proteins are responsible for the efflux of auxin out of the cell (Petrášek and Friml, 2009). The PIN proteins are considered to be highly organ- and tissue-specific. For instance, PIN1 of *Arabidopsis thaliana* is localized throughout the vascular tissues of the SAM, and in other developing organs, whereas AtPIN2, AtPIN3, AtPIN4, and AtPIN7 are found mostly in the root tip. In addition, AtPIN8 is expressed in the pollen tube (Křeček *et al.*, 2009). PIN-protein activation is dependent on phosphorylation by serine/threonine kinases, such as D6PK and PINOID (PID), and phosphorylation determines the polarity of the PIN transporters (Zourelidou *et al.*, 2014; Barbosa *et al.*, 2014). The PIN proteins are distributed throughout the plasma membrane and are directed to the membrane side where PAT is required by clathrin-based endocytosis (Dhonukshe *et al.*, 2007; Men *et al.*, 2008; Li *et al.*, 2019). Another auxin transporter family consists of the ATP-binding cassette (ABC) proteins. Some members of the ABC-B subfamily have been found to transport auxin across the plasma membrane (Titapiwatanakun and Murphy, 2009), and even increase the rate of auxin transport when they physically interact with PIN1 (Zwiewka *et al.*, 2019). Apart from membrane transporters, PAT is also dependent on cytosolic free calcium ions (Ca^+2^), as they are required for PIN1 repolarization (Li *et al.*, 2019). The calcium ions interact with PID BINDING PROTEIN 1 (PBP1) which stimulates PID phosphorylation activity on PIN1 (Benjamins *et al.*, 2003).

In alternate bearing (AB), or biennial bearing, heavy fruit load in one year (On-Crop) is followed by low fruit load in the following year (Off-Crop) (reviewed in Goldschmidt and Sadka 2020). AB is found in some cultivars of most fruit tree species, deciduous and evergreen. The intensity of the yield fluctuation differs from one species to the next, and even between cultivars of the same species. In most cases, the effect of fruit load is exerted on flower number, and it inhibits flowering induction. Flowering induction in subtropical trees, such as citrus, olive, avocado and mango, is dependent on the accumulation of a sufficient number of cold hours (Wilkie *et al.*, 2008). It is generally agreed that heavy fruit load generates an inhibitory signal (“AB signal”) that blocks flowering induction by low temperature (Goldschmidt and Sadka, 2020). Indeed, fruit removal from On-Crop trees is effective at inducing flowering, but only if it is performed in citrus no later than 1–2 months prior to the onset of the flowering-induction period (Verreynne and Lovatt, 2009; Martínez-Fuentes *et al.*, 2010; Muñoz-Fambuena *et al.*, 2011). This “memory” of the fruit load is even longer in olive, where fruit removal is only effective if performed 2 to 3 months before onset of the flowering-induction period (Dag *et al.*, 2010; Haberman *et al.*, 2017). Recent findings suggest that fruit-load memory might be controlled by epigenetic mechanisms (Agustí *et al.*, 2019).

We previously compared the transcriptomes of buds from On- and Off-Crop trees and of buds collected 1–4 weeks after complete fruit removal (defruiting) (Shalom *et al.*, 2012, 2014). Off-Crop buds and those from defruited trees showed induced expression of Ca^+2^-dependent and independent PAT genes. Accordingly, indoleacetic acid (IAA) levels were lower in Off-Crop vs. On-Crop buds, and within 1–2 weeks after defruiting, they were reduced to the level of Off-Crop buds. We suggested that fruit presence induces PAT in the stem, and this, at least in part, induces the hormone level in the bud, in accordance with the ATA theory (Bangerth, 1989). Upon fruit removal, PAT in the stem is reduced, allowing IAA release from the bud. We suggested that this might play a role in mediating the high-fruit-load signal into the bud, thereby initiating a cascade of events that lead to flowering suppression. Here, we directly investigated the effect of fruit presence on PAT in the stem and out of the bud using radiolabeled auxin in two species, citrus and olive. We show that regardless of fruit location relative to the bud—apical in citrus and basal in olive, it generates relatively high PAT in the stem, whereas fruit removal induces auxin outflow from the bud. Using immunolocalization and hormone analyses, we show that toward the flowering-induction period in citrus, IAA levels in the bud and apical meristem are higher in On-Crop trees than in defruited trees. Overall, data presented here support the notion that PAT plays a role in mediating the fruit-load signal.

## Materials and methods

### Plant material

Defruiting and sample collection were performed in a commercial orchard of ‘Murcott’ mandarin (*Citrus reticulata* x *C. sinensis*) in Nitzanim, Israel. Fruit were removed from four trees in November 2017, and a year later, in August and November 2018. Leaves, branches, and buds from defruited and fruit-bearing trees (Control) were collected in November and December of 2017 and 2018. Whole branches were brought to the laboratory on ice, before sample collection, and immediately frozen in liquid nitrogen. Then, the samples were crushed by mortar and pestle to a fine powder and kept at −80 °C. Plant material for PAT measurements in olive (*Olea europaea* cv. Barnea) was collected from the experimental farm of the ARO, Volcani Center, located in the central coastal area of Israel, where naturally On- and Off-Crop trees were located.

### Inflorescence counting

Three to four branches on each tree of more or less equal size were labeled prior to flowering time. Inflorescences and newly developed vegetative shoots were counted during the flowering peak. Flower and vegetative shoot numbers were standardized by the number of nodes present on the counted branch.

### IAA determination

Frozen bud samples were freeze-dried in a lyophilizer (Christ, Osterode am Harz, Germany) and analyzed for IAA content as described previously (Albacete *et al.*, 2008), with some modifications. Briefly, dry plant material (50 mg) was dropped into 1 ml of cold (−20 °C) extraction mixture (methanol:water, 80:20 v/v). Solids were separated by centrifugation (20,000 *g*, 15 min) and reextracted for 30 min at 4 °C in an additional 1 ml of the same extraction solution. Pooled supernatants were passed through a Sep-Pak Plus C_18_ cartridge (Sep-Pak Plus, Waters, Milford, MA, USA) to remove interfering lipids and some of the plant pigments, and then evaporated at 40 °C under vacuum to near dryness. The residue was dissolved in 1 ml methanol:water (20:80 v/v) solution using an ultrasonic bath. The dissolved samples were filtered through 13-mm diameter Millex filters with a 0.22-μm pore size nylon membrane (Millipore, Bedford, MA, USA). Filtered extract (10 μl) was injected into a U-HPLC-MS system consisting of an Accela Series U-HPLC (ThermoFisher Scientific, Waltham, MA, USA) coupled to an Exactive mass spectrometer (ThermoFisher Scientific) using a heated electrospray ionization interface. Mass spectra were obtained using Xcalibur software version 2.2 (ThermoFisher Scientific). For quantification of the plant hormones, calibration curves were constructed for each analyzed component (1, 10, 50, and 100 μg l^−1^) and corrected for 10 μg l^−1^ deuterated internal standards. Recovery percentages ranged between 92 and 95%.

### IAA immunolocalization

#### Sample preparation

IAA immunolocalization within the buds was performed essentially as described previously (Moctezuma, 1999), with some modifications. Stems were collected during the late summer from On- and Off-Crop trees and brought to the laboratory on ice. Stem sections with buds were incubated for 3 h in a solution containing 4% (v/v) 1-ethyl-3-(dimethylaminopropyl)-carbodiimide hydrochloride (EDAC), 3% (v/v) paraformaldehyde, 0.1% (v/v)? Triton X-100 in 10 mM phosphate buffer saline (PBS) pH 7.3 at room temperature, followed by overnight incubation at 4 °C in FAA solution containing 37% (v/v) formaldehyde, 10(%?) (v/v) formalin, 50% (v/v) ethanol, 5% (v/v) acetic acid. Next, the FAA solution was removed, and the samples were gradually dehydrated by 1 h incubation at room temperature in 50%, 70%, 90%, 95% ethanol, followed by 4–5 h incubation in 100% ethanol with one replacement following 1 h incubation. The ethanol was then gradually replaced with Histoclear (Bar Naor, Petach Tikva, Israel), by 1 h incubation at room temperature each with 25%, 50%, 75%, 100% Histoclear solution in ethanol. Five flakes of paraffin were then added to each bottle and incubated at 42 °C. Two to three additional paraffin flaks were added to each bottle until saturation. Next, the samples were incubated at 59 °C until the paraffin was fully dissolved, and 50% of the Histoclear solution was replaced with pure dissolved paraffin. The bottles were left open during incubation to allow Histoclear evaporation, followed by additional replacement of 50% of the solution with pure dissolved paraffin. The entire content of the bottles was then replaced with 100% paraffin, a procedure that was performed twice a day for a total of six times at 59 °C. The samples were then transferred into a ceramic mold placed on a heating plate set to 40 °C with the stem facing downwards, followed by incubation on ice for rapid solidification. Blocks of paraffin, each containing a single bud, were cut and each block was adhered to a small wood base. Samples were cut into 12-mm thick sections using a microtome (Leica RM2245, Wetzlar, Germany), and placed on charged glass slides, followed by overnight incubation at 42 °C. Section quality was evaluated by light microscopy (Leica).

#### Immunolocalization

The sections were dried on a heating plate set to 42 °C, and the paraffin was then removed by a series of washes as follows: 2 × 12 min in 100% Histoclear, 2 × 2 min in 100% ethanol, and 2 min each in 95%, 85%, 70%, 50%, 30% ethanol in PBS. Next, the sections were incubated at ambient temperature for 45 min in a blocking solution containing 0.1% (v/v) Tween-20, 1.5% (w/v) glycine, and 5% (w/v) BSA in PBS, followed by a 2-min rinse in a solution containing 10 mM PBS pH 7, 0.0088% (w/v) NaCl, 0.1% Tween-20, and 0.8% BSA (salt solution), and a 2-min rinse in a solution containing 10 mM PBS pH 7 and 0.8% BSA (BSA/PBS solution). Next, 200 μl primary antibody (rabbit anti-IAA (N1), Agrisera, Vannas, Sweden) diluted 1:1000 in PBS was added to the sections and they were covered with a cover slip. Incubation was initially performed at ambient temperature for 30 min, and then at 4 °C overnight in the dark. The sections were washed by rinsing twice for 2 min each in a solution containing 10 mM PBS pH 7, 2.9% NaCl, 0.1% Tween-20 and 0.1% BSA (high-salt solution), followed by a 2-min wash in the salt solution, and a 2-min wash in the BSA/PBS solution. Secondary antibody (anti-rabbit IgG, whole molecule, alkaline phosphatase, Sigma, St. Louis, MO, USA), 200 μl of a 1:2000 dilution in PBS, was then added to each section followed by 4 h incubation at room temperature in a moist and dark environment. The sections were then rinsed twice with salt solution followed by a ddH_2_O rinse. Detection was performed by adding 200 μl of western blue substrate (SIGMAFAST™ BCIP®/NBT = 5-bromo-4-chloro-3-indolyl phosphate/nitro blue tetrazolium, Sigma) followed by incubation at room temperature in a dark and moist environment until the reagent’s color became visible, and rinsing twice with ddH_2_O. Drying was then performed at room temperature, and the sections were covered with a cover slip and visualized by light microscopy.

### PAT

For these experiments, plant material from On-Crop citrus trees and from trees defruited 10–14 days prior to the experiment were collected during the late summer and early fall. Branches were brought to the laboratory on ice and cut to a length of about 10 cm each. Apical fruit was removed from fruit-bearing citrus branches, shortly before the beginning of the experiment. Similarly, the apical shoot of non-bearing branches of Off-Crop trees was decapitated. Plant material from olive was collected in the early summer from naturally occurring On- and Off-Crop trees. The branches were then placed vertically in 1 ml Pipetor tip containing 250 μl agar ("receiver agar"). A second agar cube of the same volume, placed in 1 ml tip, contained 5 μl (200 μM, 50 μCi) ^14^C-radiolabeled IAA (American Radiolabeled Chemicals, Saint Louis, MO, USA) with or without 5 μl of the PAT blocker 2,3,5-triiodobenzoic acid (TIBA, 1.25 mM final concentration) (Sigma-Aldrich, Rehovot, Israel). This agar cube ("donor agar") was then placed on the top of each dissected branch. For bud-labeling experiments, 1 μl (200 μM, 50 μCi) radiolabeled IAA was pipetted directly on the bud, while the donor agar either contained or did not contain 50 mM IAA. The olive experiments were set up similarly. However, first-year fruit are found in a basal position to newly developed, second-year branch-containing buds (Fig. 5A). Therefore, fruit-bearing first-year branches from On-Crop trees or non-bearing branches from Off-Crop trees were collected with second-year non-bearing branches, which were used for the experiment. The olive bud is covered with a relatively thick and impermeable bract and therefore, prior to radiolabeled IAA application, the bract was gently scraped using grade 0 sandpaper. Incubation was performed at ambient temperature in the dark for 24 or 48 h for citrus and olive, respectively. Following incubation, the branches were separated from the agar cubes and cut into three pieces of similar length and slightly chopped. The agar cube and branch pieces were placed in scintillation tubes containing 3 ml scintillation liquid (Ultima Gold, Waltham, MA, USA). When radiolabeled IAA was applied to the buds, they were separated from the branch following incubation, and washed for a few seconds in 50 μl Decon90 (diluted 1:50 in ddH_2_O) (FieldTech Solutions, Brooklyn, Australia), followed by a few seconds wash in 1 ml ddH_2_O. The Decon90 and ddH2O used for the bud washes were also placed in scintillation tubes, and they were defined as the “wash” fraction. Radioactivity was counted using a beta-scintillation counter (Packard 1900 TR, East Lyme, CT, USA). PAT was determined as the combined content of IAA, in nanomoles, present in the stem and in the receiver agar. Determination of the effect of IAA on PAT in Off-Crop trees was performed in a net house under natural conditions in October. Apical shoots of non-fruit-bearing branches were decapitated and lanolin alone or mixed with 250 ng g^−1^ IAA covered by 100 μl black Pipetor tip was applied. The lanolin was renewed every week for 1 month. Treated branches were then cut and brought to the laboratory for radiolabeled IAA application on buds, as described above.

### RNA extraction, quantification and qPCR analyses

Total RNA was extracted, treated and analyzed from approximately 0.2 g and 0.5 g of frozen bud and stem tissue, respectively, and cDNA was synthesized, as described previously (Shalom *et al.*, 2012). Primers for the analyzed genes were designed based on genomic sequences (*Citrus sinensis* annotation project, http://citrus.hzau.edu.cn/orange/) using Primer 3 software (Supplementary Table S1 at JXB online). Real-time PCR was carried out as described previously (Shalom *et al.*, 2012).

## Results

### Changes in IAA level in citrus buds of On-Crop and defruited (DEF) trees

We previously showed that fruit removal (defruiting) induces IAA levels in the citrus buds within 1 week (Shalom et al., 2014). However, defruiting in that experiment was carried out in July, much earlier than the flowering-induction period, which takes place during the winter temperature decline. Therefore, we first analyzed IAA levels during the flowering-induction period in DEF trees. During the first year, defruiting was performed in mid-November, and hormone analyses were carried out prior to fruit removal and 1 month later. Auxin was reduced by about 2-fold in the buds of DEF trees as compared to those of On-Crop trees (Fig. 1A). In the following year, defruiting was carried in either mid-August or mid-November, and IAA was analyzed in mid-November and mid-December. Defruiting in August or November resulted in about 2-fold reduced IAA levels in December (Fig. 1B). As expected, high-intensity flowering was observed when the fruit were removed in August but not when they were removed in November (Fig. 1C).

**Figure 1.**
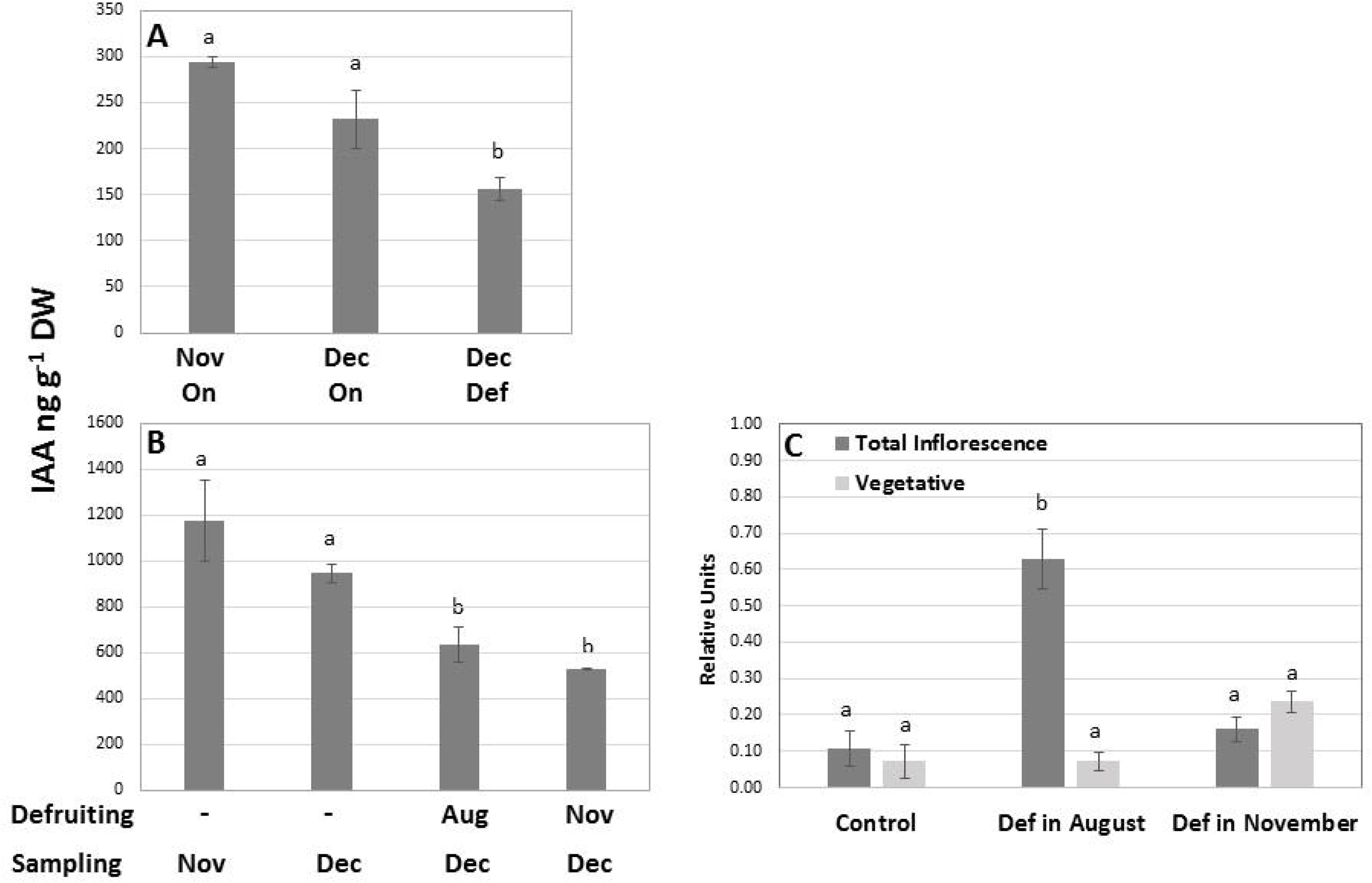
Fruit removal (defruiting) reduces auxin level in the bud during the flowering-induction period. (A) Auxin levels were determined in December 2017 in trees that were defruited (Def) in November as compared to control non-defruited trees (On), whose hormone levels were determined in November and December. (B) In 2018, tree defruiting was carried out in either August or November and hormone levels were determined in November and December, as indicated. (C) Flowers and vegetative shoots were monitored in April 2019 in On-Crop trees (Control), and in trees defruited (Def) in August 2018 or November 2018, as indicated. Relative units refer to the number of inflorescences or vegetative shoots standardized to the number of nodes per counted branch. Mean values are of three biological replicates ± SE. Different letters denote significant difference (*P* ≤ 0.05) by Student’s t-test.

Anti-IAA antibodies were used to label IAA in the buds of On- and Off-Crop trees. IAA labeling was detected in the apical meristem of On-Crop buds, but was barely detectable in that of Off-Crop buds (Fig. 2).

**Figure 2.**
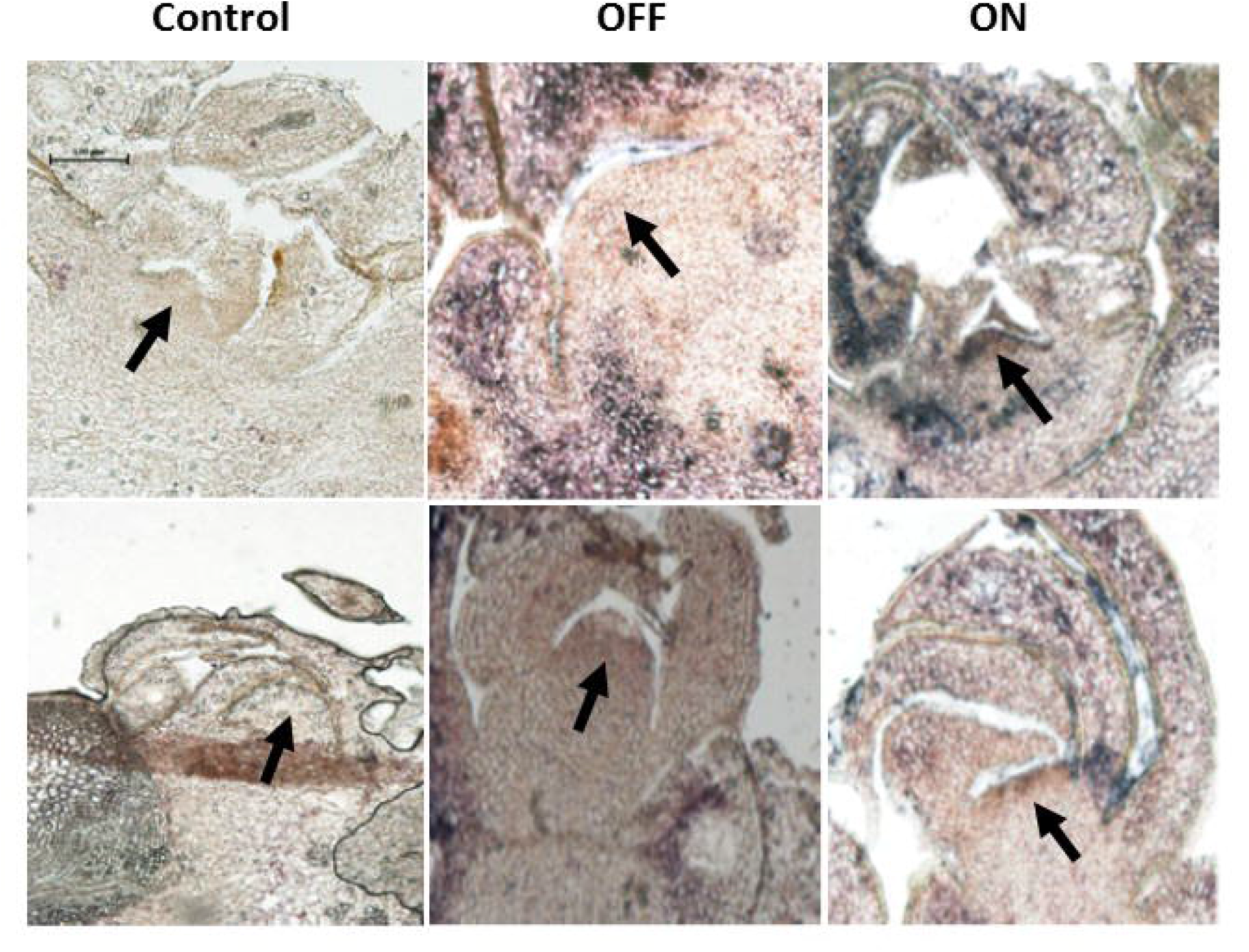
Auxin levels are lower in the shoot apical meristem of On-Crop vs. Off-Crop buds. Immunolocalization of IAA in longitudinal sections of two buds sampled from On-Crop trees (ON) or defruited trees (OFF), as indicated. Arrow indicates the apical meristem. Control refers to buds immunolabeled with primary antibodies only (top control photo) or secondary antibodies only (bottom control photo).

### Fruit load affects PAT in citrus stems and hormone release from the buds

Our previous work (Shalom et al., 2014) and the above data suggested that PAT is induced in the stem by the presence of fruit and that hormone release from the bud is reduced. We examined this hypothesis using radiolabeled auxin. ^14^C-IAA was applied in donor agar placed on top of a young decapitated stem taken from a DEF tree or a fruit-bearing branch of an On-Crop tree (the fruit was removed just prior to the experiment) (Fig. 3A)(. PAT was determined by counting the radiolabel in the stem and in the receiver agar. Auxin transport was almost 10-fold higher in the On-Crop stem compared to the DEF stem (Fig. 3B). Addition of the PAT blocker TIBA to the donor agar of the On-Crop branch, or applying the radiolabeled auxin to the donor agar of a stem incubated upside-down (reversed), almost abolished PAT.

**Figure 3.**
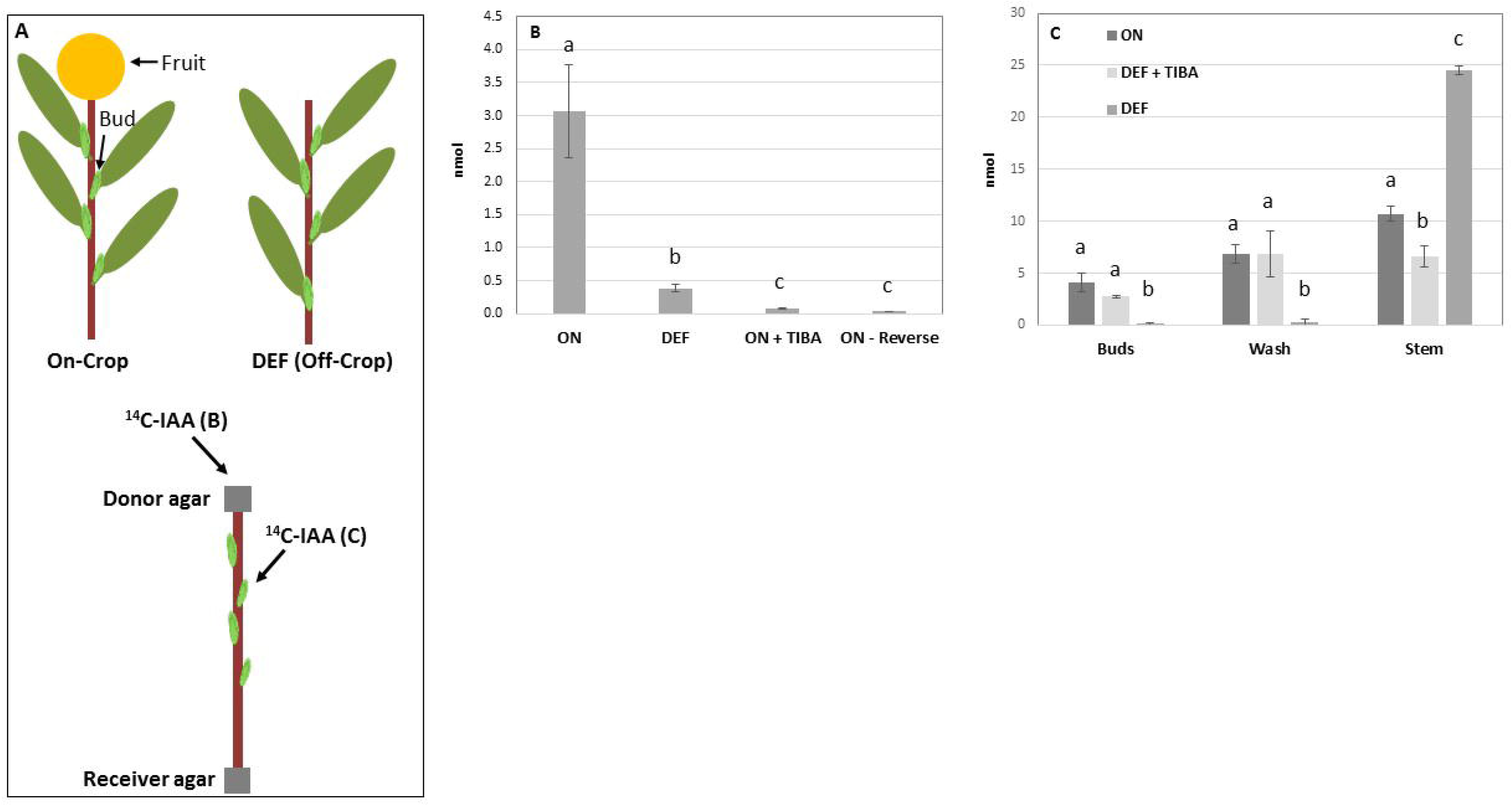
Polar auxin transport in citrus stem is affected by fruit load while defruiting allows auxin release from the bud. (A) Radiolabeled auxin (^14^C-IAA) was applied in donor agar placed on top of a branch sampled from a defruited tree (DEF) or an On-Crop tree from which the fruit was removed just prior to the experiment, or a branch from an On-Crop tree incubated upside-down, as illustrated schematically (^14^C-IAA (B)). (B) Polar auxin transport (PAT) was determined by counting the radiolabeled IAA in the shoot and in the receiver agar following 24 h of incubation in the shoots of On-Crop trees (ON), defruited trees (DEF), On-Crop trees with application of TIBA along with the radiolabeled IAA (ON + TIBA), and On-Crop trees incubated upside-down (ON-Reverse). (C) ^14^C-IAA was applied to a bud on a branch sampled from a defruited tree (DEF) or an On-Crop tree from which fruit was removed just prior to the experiment (ON), as illustrated schematically (A, ^14^C-IAA (C)). PAT was determined by counting the radiolabeled IAA in the stem and in the receiver agar (Stem), as well as in the bud (Buds) and in the bud wash solution (Wash) following 24 h of incubation; DEF + TIBA, defruited trees to which TIBA was applied along with the radiolabeled IAA. Mean values are of four biological replicates ± SE. Different letters denote significant difference (*P* ≤ 0.05) by Student’s t-test.

Next, we examined the effect of fruit presence on auxin release from a lateral bud (Fig. 3C). Following ^14^C-IAA application to the bud and incubation, the bud was dissected and washed thoroughly to remove non-penetrated auxin. Radiolabeled auxin was greatly reduced in the bud of the DEF tree as compared to that of the On-Crop tree or the DEF bud to which TIBA was applied (Fig. 3C). A ca. 2-to 3-fold increase in radiolabeled auxin was detected in the DEF stem compared to the On-Crop stem and the DEF stem with applied TIBA. Interestingly, auxin penetration into the bud was remarkably higher in the DEF bud as compared to the On-Crop bud or the DEF bud with TIBA, as evidenced by the amount of radiolabel remaining in the wash solution.

To determine whether fruit presence can be replaced by IAA, non-radioactive hormone was applied to the decapitated stem of an Off-Crop tree, and radiolabeled IAA was applied to the bud (Fig. 4A). ^14^C-IAA content in the bud of the Off-Crop tree with cold hormone was similar to that of the On-Crop tree, and was reduced by about 2-to 3-fold in the bud of the Off-Crop tree with no applied cold hormone (Fig. 4B). In parallel, IAA content in the stem of the Off-Crop tree with applied cold hormone was similar to that of the On-Crop tree, and was induced by about 2-fold in the bud of the Off-Crop tree. ^14^C-IAA penetration into the bud of the Off-Crop tree with cold hormone was similar to that of the On-Crop tree, as evidenced by the radiolabel counts in the wash solution.

**Figure 4.**
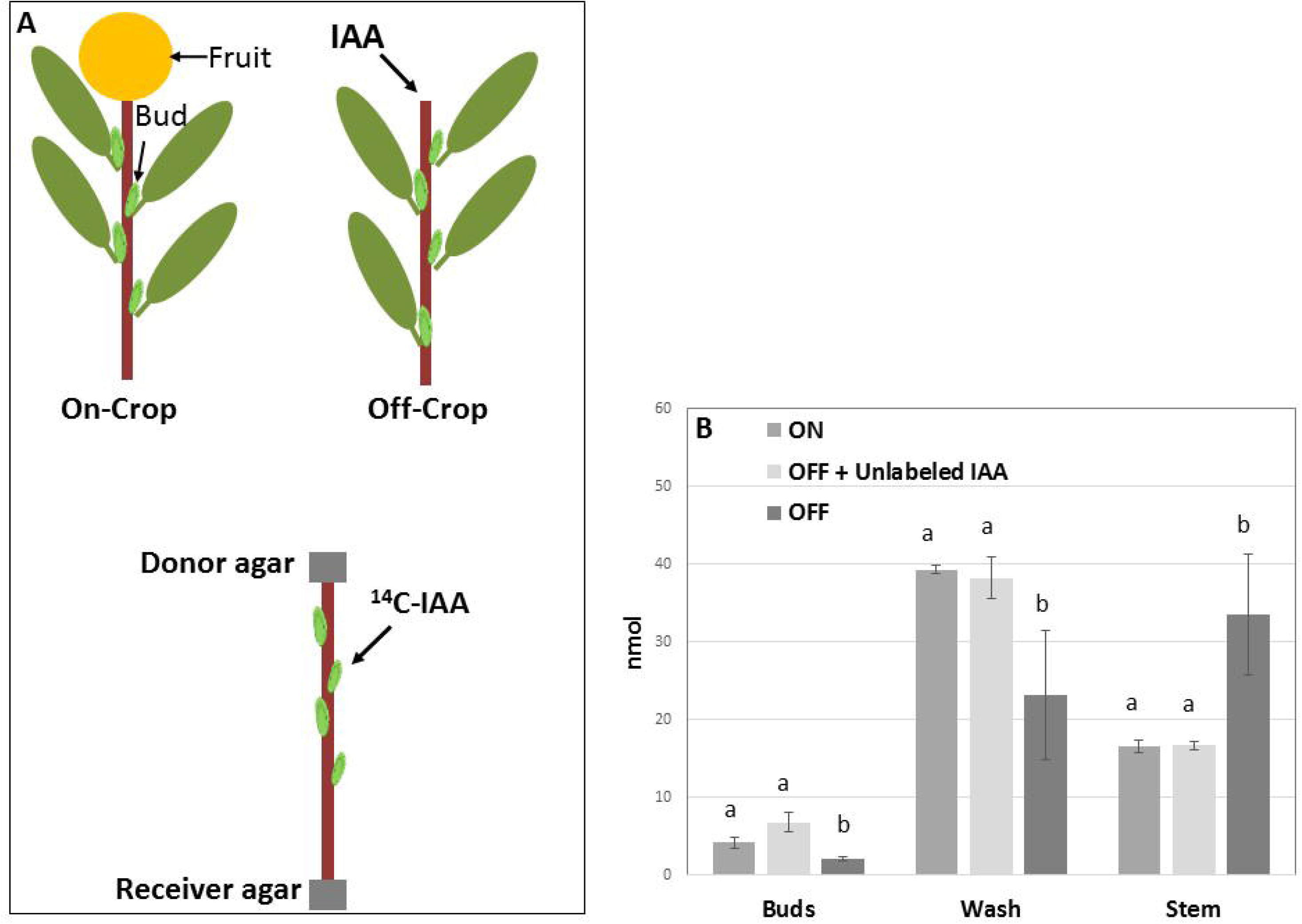
Auxin can replace the fruit when applied to branches of Off-Crop trees. Non-radiolabeled IAA was applied to branches sampled from Off-Crop trees prior to application of radiolabeled IAA (^14^C-IAA) to the buds of those branches and to branches sampled from On-Crop trees. (A) Schematic representation of the experiment, indicating the site of application of the radiolabeled IAA (^14^C-IAA). (B) Polar auxin transport (PAT) was determined by counting the radiolabeled IAA in the stem and in the receiver agar (Stem), as well as in the bud (Buds) and in the bud wash solution (Wash), following 24 h of incubation, in samples of On-Crop trees (ON) where fruit was removed just prior to the experiment, and Off-Crop trees with application of non-radiolabeled IAA (OFF + Unlabeled IAA) and Off-Crop trees (OFF). Mean values are of four biological replicates ± SE. Different letters denote significant difference (*P* ≤ 0.05) by Student’s t-test.

**Figure 5.**
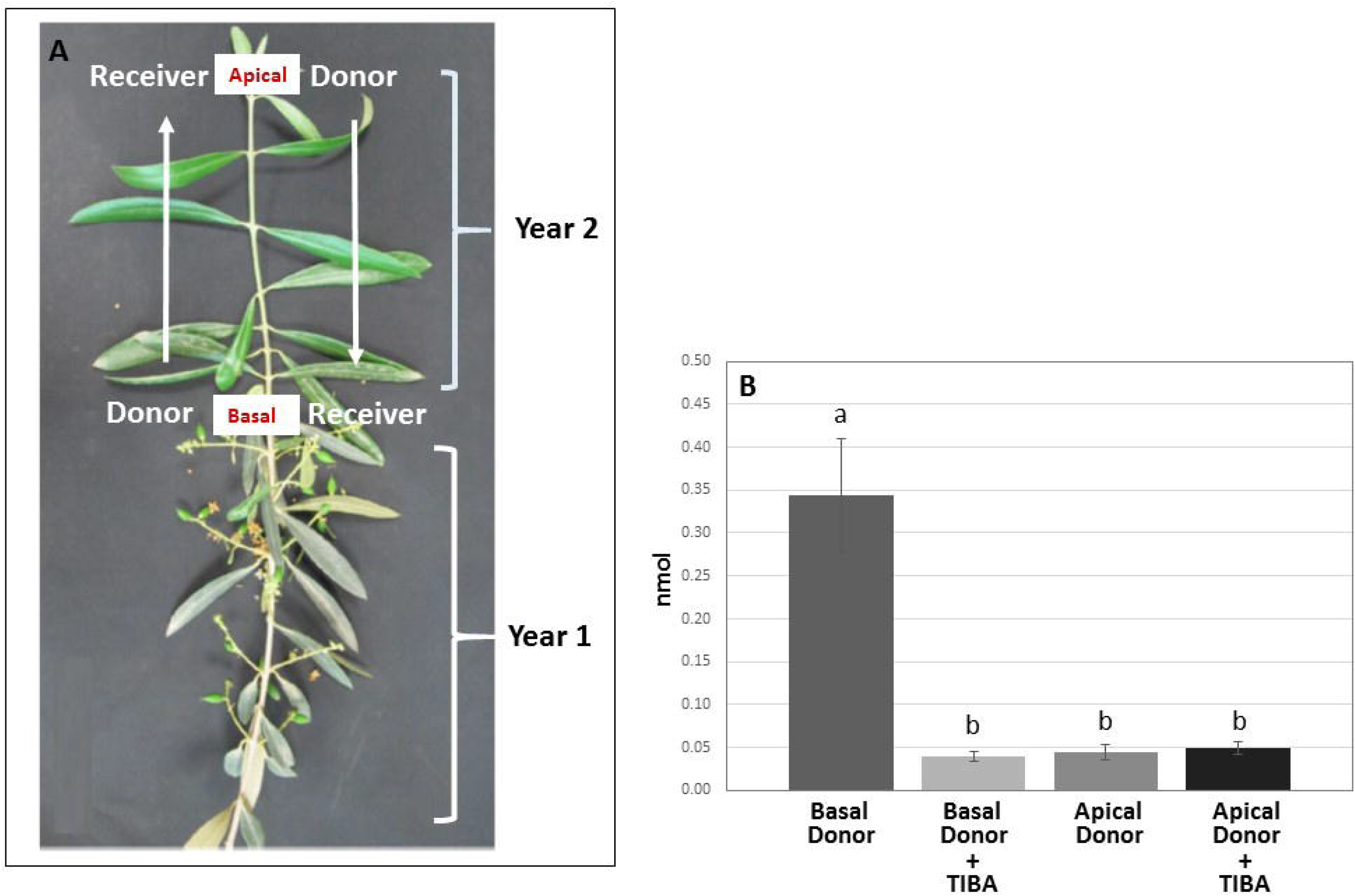
Basal to apical polar auxin transport in olive. (A) In olive, first-year fruit are basal to the second-year branch. To determine the direction of polar auxin transport (PAT), radiolabeled IAA was applied either basally or apically through donor agar to a second-year branch. (B) PAT in the second-year branch was detected when ^14^C-IAA was applied basally but not apically, and it was inhibited by the addition of TIBA. Means values are of four biological replicates ± SE. Different letters denote significant difference (*P* ≤ 0.05) by Student’s t-test.

### Fruit presence at the basal position affects stem PAT and auxin release in olive

The effect of fruit load on PAT was examined in olive, where the first-year fruit-bearing stem inhibits flowering induction in the second-year stem, so the fruit is in a basal position relative to the bud (Fig. 5A). To validate PAT direction, ^14^C-IAA was applied either apically or basally to the second-year shoot of an On-Crop tree. Auxin levels were about 10-fold higher when it was applied basally as compared to apically, or with apical/basal application of TIBA (Fig. 5B), demonstrating that the direction of PAT is basal to apical. The effect of fruit presence on PAT was then determined by applying ^14^C-IAA to agar placed at the basal position of stems from On- and Off-Crop trees (Fig. 6A). Radiolabeled auxin content was reduced by about 4-fold in the stem of the Off-Crop tree, as compared to its content in the stem of the On-Crop tree (Fig. 6B). Application of TIBA reduced PAT in the On-Crop stem about 2-fold, and incubating the On-Crop stem upside-down (reverse) reduced auxin content by about 3-fold. Overall, although consistent with the citrus data, the effect of fruit presence was less dramatic in olive. Next, the effect of fruit presence on IAA bud content was examined by applying ^14^C-IAA to On-Crop and Off-Crop buds. A ca. 2-fold decrease in IAA level in the bud and a similar increase in the stem were evident in Off-Crop trees compared to On-Crop trees, with TIBA abolishing the effect (Fig. 6C).

**Figure 6.**
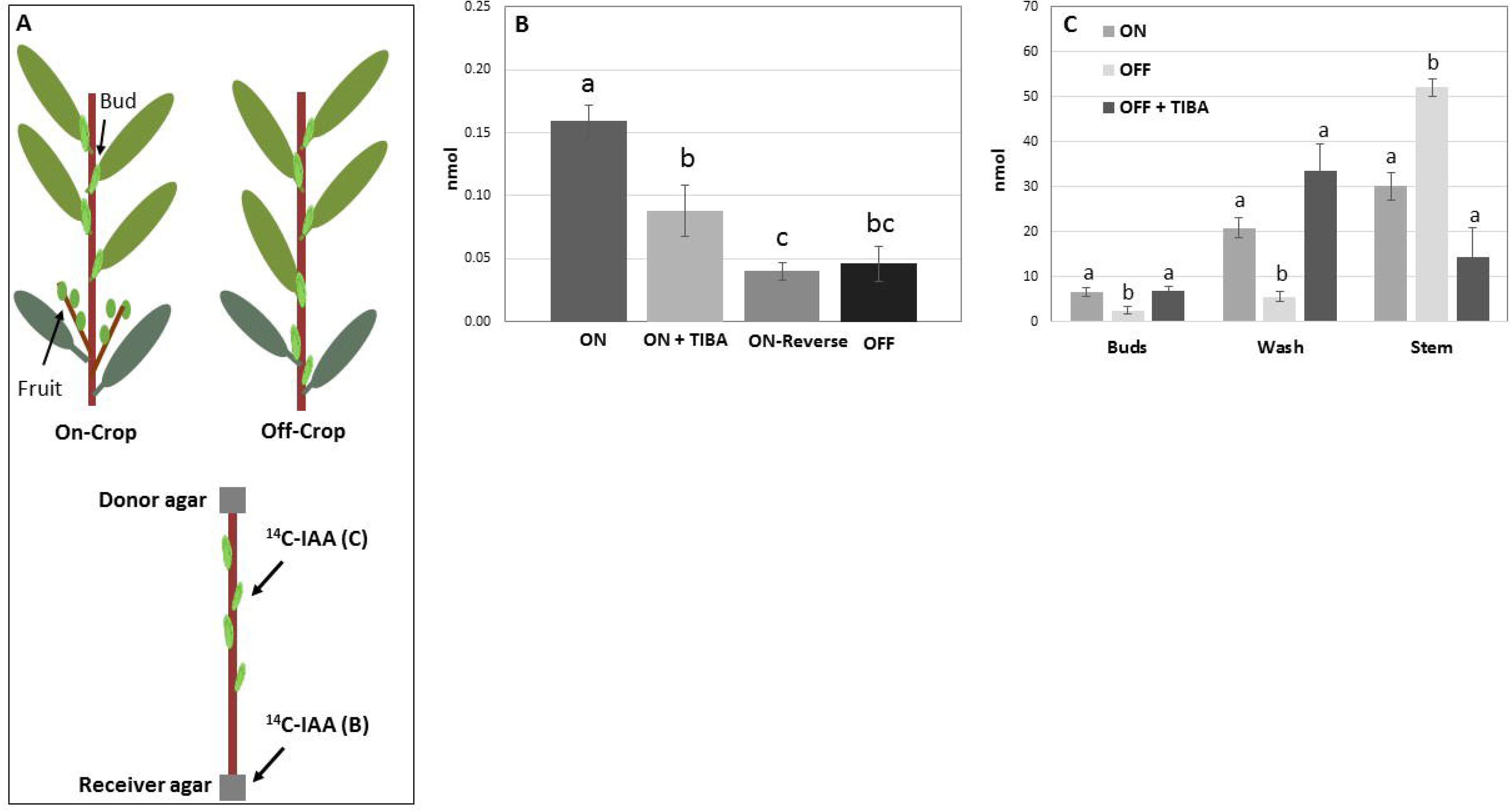
Polar auxin transport direction is affected by fruit load in olive, while fruit absence induces hormone release from the bud. (A) Radiolabeled auxin (^14^C-IAA) was applied in donor agar placed at the bottom of a second-year branch sampled from an Off-Crop tree or at the bottom of a branch sampled from an On-Crop tree, as illustrated schematically (^14^C-IAA (B)). (B) Polar auxin transport (PAT) was determined by counting the radiolabeled IAA in the shoot and in the receiver agar following 48 h of incubation of On-Crop trees (ON), Off-Crop trees (OFF), On-Crop trees with application of TIBA + radiolabeled IAA (ON + TIBA), and On-Crop trees with the shoot incubated upside-down (ON-Reverse). (C) Radiolabeled auxin (^14^C-IAA) was applied to a bud on a branch sampled from an Off-Crop tree or from an On-Crop tree, as illustrated schematically (A, ^14^C-IAA (C)). PAT was determined by counting the radiolabeled IAA in the stem and in the receiver agar (Stem), as well as in the bud (Buds) and in the bud wash solution (Wash) following 48 h of incubation of On-Crop trees (ON), Off-Crop trees (OFF), and Off-Crop trees with TIBA application along with the radiolabeled IAA (OFF + TIBA). Mean values are of four biological replicates ± SE. Different letters denote significant difference (*P* ≤ 0.05) by Student’s t-test.

### Changes in auxin-related genes upon fruit removal

We previously published the transcriptomes of On and Off-Crop buds, and those of buds 1, 2 and 4 weeks after defruiting (Shalom al., 2014). Supplementary Table S2 presents the ratios of transcript levels between On- and Off-Crop buds and between On-Crop and DEF buds of genes related IAA metabolism, transport and signaling. Most of the genes associated with IAA conjugation were induced in On-Crop buds as compared to Off-Crop buds or buds of DEF trees, in association with the induced IAA level. LAX2, which plays a role in auxin transport, was induced following defruiting, in association with IAA release from the bud, but PIN5 and PIN6 showed the opposite trend: they were higher in On-Crop buds. Notably, some PIN proteins might play a role in PAT outside the bud, and some in IAA movement within the bud, which might explain these results (Křeček *et al.*, 2009). For IAA signaling/response, AUX-IAA genes showed higher expression in On-Crop buds, while expression of SUAR genes was reduced, but this might reflect the normal auxin response in which some genes are induced and some are reduced. Interestingly, one detected IAA-biosynthesis gene, YUCCA10 and one auxin receptor, TIR1, were induced in Off-Crop buds and/or following defruiting.

The above results demonstrate changes in the bud transcriptome following summer defruiting, but not during the flowering-induction period. We therefore examined changes in gene expression during the fall and winter in buds and stems of trees that had been defruited in July. YUCCA10 showed induced expression in On-Crop buds compared to DEF buds from July to September and a decline thereafter (Fig. 7A); this pattern of induction and then decline was also seen in the stem, but for both On-Crop and DEF trees (Fig. 7B). A similar pattern was detected for IAA methyltransferase, which was higher in both buds and stems of On-Crop trees as compared to those of DEF trees in September, with bud transcript levels showing slight induction from December to January (Fig. 7C and D). Overall, GH3.5 transcript levels decreased from July to January in both tissues with no obvious difference between buds of On-Crop and DEF trees. However, from December to January, GH3.5 expression was induced in stems of On-Crop trees, but not in those of DEF trees (Fig. 7E and F). The transcript levels of IAA UDP-glycosyltransferase and IBA (indole-3-butyric acid) UDP-glycosyltransferase genes did not show remarkable differences between On-Crop buds and stems and those of DEF trees (Supplementary Fig. S1). Transcript levels of the three analyzed auxin transporter genes, PIN1, PIN3 and ABCB19, displayed overall similar pattern in the buds, with all of them showing induction from July to September and from December to January (Fig. 8A, C and E). Transcript levels of all three genes were lower in On-Crop buds than in DEF buds. In the stems, PIN1 transcript levels decreased from September to January, with slightly but significantly higher transcript levels in DEF vs. On-Crop trees (Fig. 8B). PIN3 transcript fluctuated with increases in September and January, and slightly higher transcript levels in DEF trees than in On-Crop ones. ABCB19 showed an overall similar pattern, with an increase in transcript levels from July to September only in stems of DEF trees. We also tested the expression of genes known to phosphorylate PIN proteins, PID and D6PK1 and 2 (Supplementary Fig. S2), but their transcript levels were similar in tissues of On-Crop and DEF buds, in agreement with reports of these proteins’ post-transcriptional regulation (Willige *et al.*, 2013). Nevertheless, PID showed lower levels in buds of DEF trees as compared to On-Crop buds in September.

**Figure 7.**
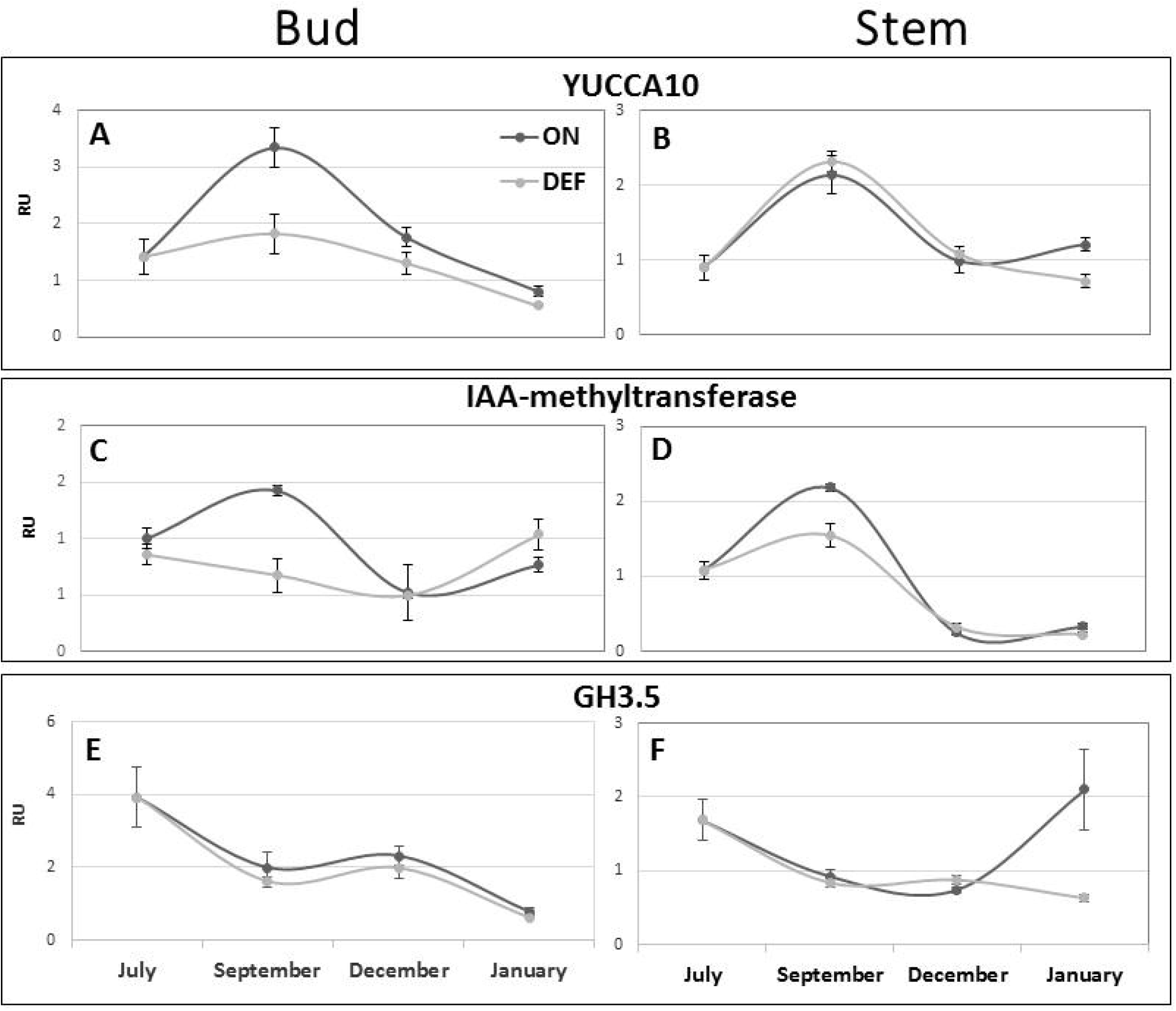
Fruit load effect on transcript levels of IAA-metabolizing genes. The indicated genes (for their ID, see Supplementary Table S1 were analyzed during the indicated months in buds (A,C,E) and stems (B,D,F) of On-Crop trees (ON) and in tissues of trees that were defruited in July (DEF). Mean values of three biological replicates ± SE. RU, relative units.

**Figure 8.**
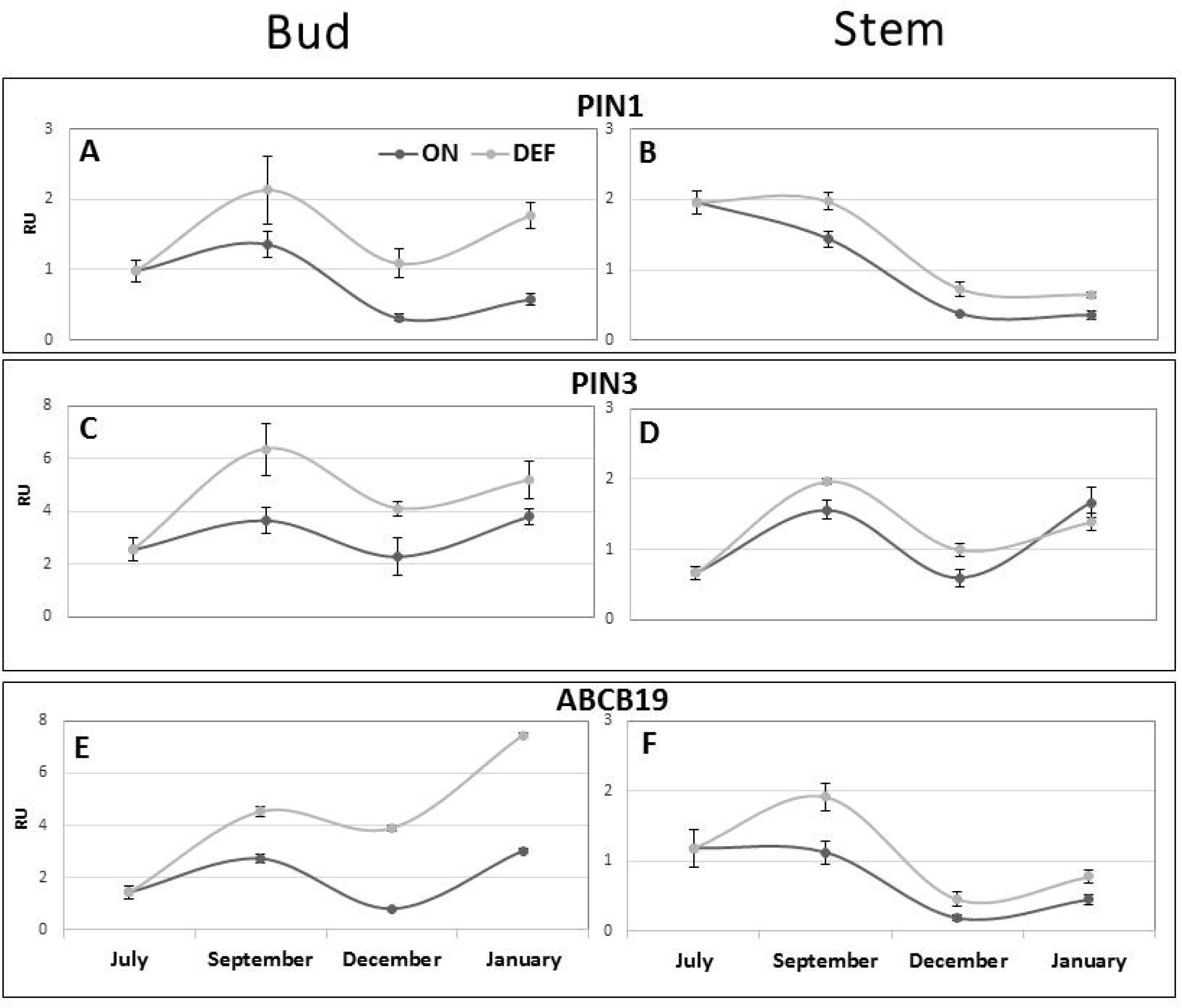
Fruit load effect on transcript levels of IAA transporters. The indicated genes (for their ID, see Supplementary Table S1 were analyzed during the indicated months in buds (A,C,E) or stems (B,D,F) of On-Crop trees (ON) and in tissues of trees that were defruited in July (DEF). Mean values of three biological replicates ± SE. RU, relative units.

## Discussion

The possible involvement of auxin in mediating a yield effect on the return bloom was first tested by examining its levels in the bud following fruit removal. In our previous report, defruiting in July reduced auxin levels in the bud in a relatively short time, 1 to 4 weeks (Shalom *et al.*, 2014). However, in that report, we did not examine the effect of fruit removal during flowering induction. Here, during 2 years, fruit removal in August or November reduced auxin levels in December (Fig. 1), providing support for the notion that the hormone plays a role in the AB signal. Furthermore, immunolocalization of auxin in the bud showed its presence in the apical meristem, and it was higher in On-Crop buds than in Off-Crop buds. Auxin localization in the SAM has been widely reported; its biosynthesis and accumulation in the SAM has been associated with self-organization of the meristem and organogenesis (Vernoux *et al.*, 2010; Sassi and Vernoux, 2013). This was clearly demonstrated during Arabidopsis embryogenesis, where PAT from the SAM controlled some aspects of embryo development, including cotyledon development, and apical–basal axis formation (Robert *et al.*, 2013; Caragea and Berleth, 2017). Our data indicate that auxin transmits the heavy-fruit-load signal to the SAM, thus playing a role in determining its fate—vegetative growth, flowering or developmental arrest—and providing a unique case of auxin action in the SAM.

Auxin distribution in plant organs is dependent on its polar transport. Therefore, changes in auxin levels in the bud and stem should be associated with PAT. As already noted, according to the ATA theory, the fruit, as a strong sink, usually dominates the bud regardless of its relative position—apical, basal or other (Bangerth, 1989). In ‘Murcott’ mandarin, most of the fruit are apical to the buds. However, in olive, the first-year On-Crop shoot continues to grow during the second year, and becomes an Off-Crop shoot, so the fruit are basal to the second-year buds. By monitoring auxin transport in these two species, we showed that the direction of PAT in the stem originates from the fruit, regardless of its position relative to the bud. We did not monitor PIN activities and localization in the stem. However, the radiolabeled IAA experiments, which included use of the PAT blocker TIBA and measuring auxin transport in “reversed” stems, demonstrated the existence of relatively strong PAT in the stem when fruit was present, and reduced PAT upon fruit removal, as shown in other studies using radiolabeled auxin (Kaldewey, 1984; Chabikwa *et al.*, 2019). In addition, fruit presence could be replaced by external application of IAA to stems sampled from Off-Crop trees, at least in citrus. Furthermore, PAT strength in the stem was strongly associated with IAA release from the bud, as suggested by the ATA theory (Bangerth, 1989; Callejas and Bangerth., 1998). A 2-to 3-fold increase in the transcript levels of three IAA transporters, PIN1, PIN2 and ABCB19, in the buds of DEF trees as compared to On-Crop buds provided support for these conclusions (Fig. 8A,C,E). Although major regulation of PIN is thought to be at the post-transcriptional level, these results were in accordance with the induction of PIN expression in other systems (Vieten *et al.*, 2005; Chabikwa *et al.*, 2019). Moreover, the induction in transcript levels of PIN1 and ABCB19 coincided with the onset of flowering induction. Interestingly, in the stem, transcript levels of all three genes were slightly higher in DEF trees than in On-Crop ones; nevertheless, PIN1 and ABCB19 seemed to demonstrate an overall decrease from September toward the induction period. Further support was provided by the induced expression of YUCCA10 and IAA methyltransferase in September in the On-Crop buds, which coincides with overall higher levels of IAA in these buds as compared to those of DEF trees (Chabikwa *et al.*, 2019).

An open question remains regarding the origin of IAA in the stem. It was previously hypothesized that fruit, and specifically seeds, are the source for auxin production (Callejas and Bangerth, 1998), a hypothesis which was recently supported by findings in Arabidopsis (Ware *et al.*, 2020). Both the citrus and olive cultivars used here have seedy fruit. However, AB also occurs in seedless cultivars (Monerri *et al.*, 2011), which raises the question of the source of IAA in those cultivars. Nevertheless, some biosynthesis may occur in the stem itself, as transcript levels of YUCCA10 and IAA methyl transferase were induced from July toward September, with the On-Crop stem showing slightly higher levels of IAA methyl transferase than stems of DEF trees. Transcript levels of GH3.5 were also induced during flowering induction from December to January in On-Crop stems, whereas no such induction was evident in the stems of DEF trees.

Although fruit removal in November reduced auxin levels, return bloom did not occur (Fig. 1C), in accordance with previous findings that citrus defruiting is effective only until September–October (Verreynne and Lovatt, 2009; Martínez-Fuentes *et al.*, 2010; Muñoz-Fambuena *et al.*, 2011). This suggested that although bud levels of auxin might play a role in mediating the fruit-load effect, this is not the sole factor affecting flowering, and other components or mechanisms are likely to be involved. Indeed, it has been generally agreed that auxin is not involved in flowering induction, but only at later stages of floral organ termination and formation (Yamaguchi *et al.*, 2018). So the question could be raised as to the cascade of events that lead to inhibition of flowering induction. Fruit presence is thought to generate a “memory” which can be reversed by fruit removal until a certain time prior to the onset of the flowering-induction period, usually 1 month in citrus. However, in olive, the “memory” of fruit load seems to last longer, as fruit removal is effective at inducing a return bloom only if it is performed 2–3 months prior to flowering induction (Dag *et al.*, 2010; Haberman *et al.*, 2017). Solid data exist on the involvement of gibberellin (GA) in flowering control in Arabidopsis and fruit trees. Whereas in Arabidopsis, GA promotes the transition of the apical meristem to a reproductive one, in fruit trees, the hormone represses flowering, in association with repressed expression of flowering-control genes such as *FT*, *AP1* and *LFY* (González-Rossia *et al.*, 2007; Monerri *et al.*, 2011; Muñoz-Fambuena *et al.*, 2012; Goldberg-Moeller *et al.*, 2013; Elsysy and Hirst, 2019). Solid data also exist on the effect of auxin on GA biosynthesis during fruit set (De Jong *et al.*, 2009; Hu *et al.*, 2018), and during seed and fruit maturation (Wolbang and Ross, 2001; Frigerio *et al.*, 2006; Desgagné-Penix and Sponsel, 2008). We therefore hypothesize that induced levels of IAA in the bud, resulting from the heavy fruit load, lead to induced levels of GA, which acts as a flowering inhibitor. Furthermore, as long as GA has not been induced, fruit removal is effective at inducing a return bloom. According to this hypothesis, GA induction in olive buds takes place earlier than in citrus, explaining the relatively long period of fruit-load “memory” in olive as compared to citrus. Fruit-load “memory” has been suggested to be associated with epigenetic regulation of citrus *MADS19*, an *LOWERING LOCUS C (FLC)* ortholog, which suppresses the expression of *FLOWERING LOCUS T* (*FT*) in the vernalization/cold pathway (Agustí *et al.*, 2019). Therefore, along with the above hypothesis, it is worth investigating whether GA also affects FLC expression. Regardless of the nature of the flowering-inhibition signal, transcript levels of a few of the analyzed genes involved in IAA synthesis, transport and conjugation (YUCCA10, IAA methyltransferase, PIN1, PIN3, PID, D6PKL) peaked in September, in association with the onset of the final period during which defruiting was effective at inducing a return bloom, and in association with induction of the flowering-inhibition signal.

In summary, despite the lack of transgenic approaches for use in model or easily transformed plants to study PAT, such as *DR5* promotor and PIN–GFP fusion proteins (Mravec *et al.*, 2008), we demonstrate a connection between fruit load and auxin homeostasis and PAT within the bud and stem. However, the role of auxin in fruit-load-associated meristematic fate requires further investigation: first, at the mechanistic level, i.e., to demonstrate the effect of fruit load on PIN expression and/or its distribution in the stem and bud, and second, to demonstrate more directly the effect of the hormone on bud fate: flowering or vegetative. These two directions are currently being investigated in our laboratory.

## ABBREVIATIONS

AB: alternate bearing
ATA: auxin transport auto-inhibition
IAA: indoleacetic acid
PAT: polar auxin transport
SAM: shoot apical meristem

## Acknowledgements

This research was supported by grant number 20-10-0058 from the Research Fund of Chief Scientist of the Ministry of Agriculture and Rural Development, Israel

## Data availability statement

All the data, including original real time PCR and PAT raw data, is available upon request.

## Author contributions

DH, LS, YL, LS, IK and MM collected the plant material and/or performed the experiments, AAAM and RMR performed hormonal analyses, AS and DH wrote the manuscript

## Supplementary material

**Table S1. Primers used in this work for real-time PCR. Table S2. Transcript levels of genes of IAA metabolism, conjugation, transport and signaling.** Ratios of transcript levels between On- and Off-Crop buds (ON and OFF, respectively) or between On-Crop buds and those of defruited trees (DEF) at the indicated times following defruiting; w, week. Data taken from Shalom et al. (2014).

**Figure S1. Fruit load effect on transcript levels of IAA-metabolizing genes.** The indicated genes (for their ID, see Table S1 were analyzed during the indicated months in buds (A,C) or stems (B,D) of On-Crop trees (ON) and trees that were defruited in July (DEF). Mean values of three biological replicates ± SE. RU, relative units.

**Figure S2. Fruit load effect on transcript levels of IAA transporters.** The indicated genes (for their ID, see Table S1 were analyzed during the indicated months in buds (A,C) or stems (B,D,E) of On-Crop trees (ON) and trees that were defruited in July (DEF). Mean values of three biological replicates ± SE. RU, relative units.

